# Repurposing of the antibiotic nitroxoline for the treatment of mpox

**DOI:** 10.1101/2022.12.29.522228

**Authors:** Denisa Bojkova, Nadja Zöller, Manuela Tietgen, Katja Steinhorst, Marco Bechtel, Tamara Rothenburger, Joshua D. Kandler, Julia Schneider, Victor M. Corman, Sandra Ciesek, Holger F. Rabenau, Mark N. Wass, Stefan Kippenberger, Stephan Göttig, Martin Michaelis, Jindrich Cinatl

## Abstract

The antiviral drugs tecovirimat, brincidofovir, and cidofovir are considered for mpox (monkeypox) treatment despite a lack of clinical evidence. Moreover, their use is affected by toxic side-effects (brincidofovir, cidofovir), limited availability (tecovirimat), and potentially by resistance formation. Hence, additional, readily available drugs are needed. Here, therapeutic concentrations of nitroxoline, a hydroxyquinoline antibiotic with a favourable safety profile in humans, inhibited the replication of 12 mpox virus isolates from the current outbreak in primary cultures of human keratinocytes and fibroblasts and a skin explant model by interference with host cell signalling. Tecovirimat, but not nitroxoline, treatment resulted in rapid resistance development. Nitroxoline remained effective against the tecovirimat-resistant strain and increased the anti-mpox virus activity of tecovirimat and brincidofovir. Moreover, nitroxoline inhibited bacterial and viral pathogens that are often co-transmitted with mpox. In conclusion, nitroxoline is a repurposing candidate for the treatment of mpox due to both antiviral and antimicrobial activity.

## Introduction

Two clades of mpox (previously known as monkeypox) virus, a member of the genus *Orthopoxvirus* in the family *Poxviridae*, caused until recently only limited zoonotic outbreaks in Africa [Elsayed et al., 2022; Gessain et al., 2022; Huang et al., 2022; Mitjà et al., 2022; Rabaan et al., 2022]. Currently, mpox viruses considered as clade IIB (occasionally also as clade III, consensus on the nomenclature is still developing) are spreading for the first time by sustained human-to-human transmission outside of Africa [Elsayed et al., 2022; Gessain et al., 2022; Huang et al., 2022; Mitjà et al., 2022; Rabaan et al., 2022]. This ongoing outbreak was classified as a ‘Public Health Emergency of International Concern’ by the WHO on 23^rd^ July 2022 [Elsayed et al., 2022; Gessain et al., 2022; Huang et al., 2022; Mitjà et al., 2022; Rabaan et al., 2022] and has at the time of writing (29th December 2022) affected at least 110 countries, accounting for 83,539 documented cases and at least 72 deaths [CDC, 2022].

About 10% of patients require hospital treatment in the current global outbreak, mainly due to pain and bacterial superinfections [Fink et al., 2022; Gessain et al., 2022; Girometti et al., 2022; Patel et al., 2022; Thornhill et al., 2022]. This is in contrast to the disease severity observed in the endemic mpox areas in Africa, in which mpox outbreaks are associated with mortality rates of up to 12% [Mitjà et al., 2022; Qiu et al., 2022; Singh et al., 2022]

Three antiviral drugs (tecovirimat (ST-246), brincidofovir (CMX001), cidofovir) are mainly considered for mpox treatment, although they have not undergone clinical testing for mpox treatment [DeLaurentis et al., 2022; Elsayed et al., 2022; Gessain et al., 2022; Huang et al., 2022; Mitjà et al., 2022; Rabaan et al., 2022]. Despite differences in the clinical presentation of the current mpox outbreak compared to previous ones [Gessain et al., 2022; Girometti et al., 2022; Hoffmann et al., 2022; Huang & Wang, 2022; Iñigo Martínez et al., 2022], recent findings indicated that these three drugs are still effective against the currently circulating mpox viruses in therapeutically achievable concentrations [Frenois-Veyrat et al., 2022; Warner et al., 2022; Bojkova et al., 2022].

Notably, the use of cidofovir and brincidofovir is associated with severe, therapy-limiting side effects [Adler et al., 2022; Gessain et al., 2022]. Moreover, the availability of tecovirimat is limited and may be affected by resistance formation [DeLaurentis et al., 2022; Gessain et al., 2022; Johri et al., 2022; Pfäfflin et al., 2022]. Hence, additional effective and readily available drugs are needed for the treatment of mpox.

Here, we investigated the antibiotic nitroxoline, which is used as a first-line therapy for uncomplicated urinary tract infections [Naber et al., 2014; Dobrindt et al., 2021; Wykowski et al., 2022], for activity against mpox viruses. Nitroxoline is known to inhibit the PI3K/AKT/mTOR and Raf/MEK/ERK signalling pathways [Chang et al., 2015; Xu et al., 2019; Palicelli et al., 2021], which are known to be critically involved in orthopoxvirus replication [Kindrachuk et al., 2012; Beerli et al., 2019; Peng et al., 2020]. As an antibiotic, nitroxoline also has the potential to target sexually transmitted bacteria that are commonly co-transmitted with mpox virus during the current outbreak and can aggravate mpox disease [Girometti et al., 2022; Hoffmann et al., 2022; Huang & Wang, 2022; Iñigo Martínez et al., 2022].

## Results

### Effects of nitroxoline on mpox virus replication

The effect of the 8-hydroxyquinoline derivative nitroxoline (Figure 1A) was determined on the replication of 12 mpox virus isolates (Suppl. Table 1) from the current global outbreak cultured in primary human foreskin fibroblasts (HFF) and primary human foreskin keratinocytes (HFK) as previously described [Bojkova et al., 2022].

**Figure 1.**
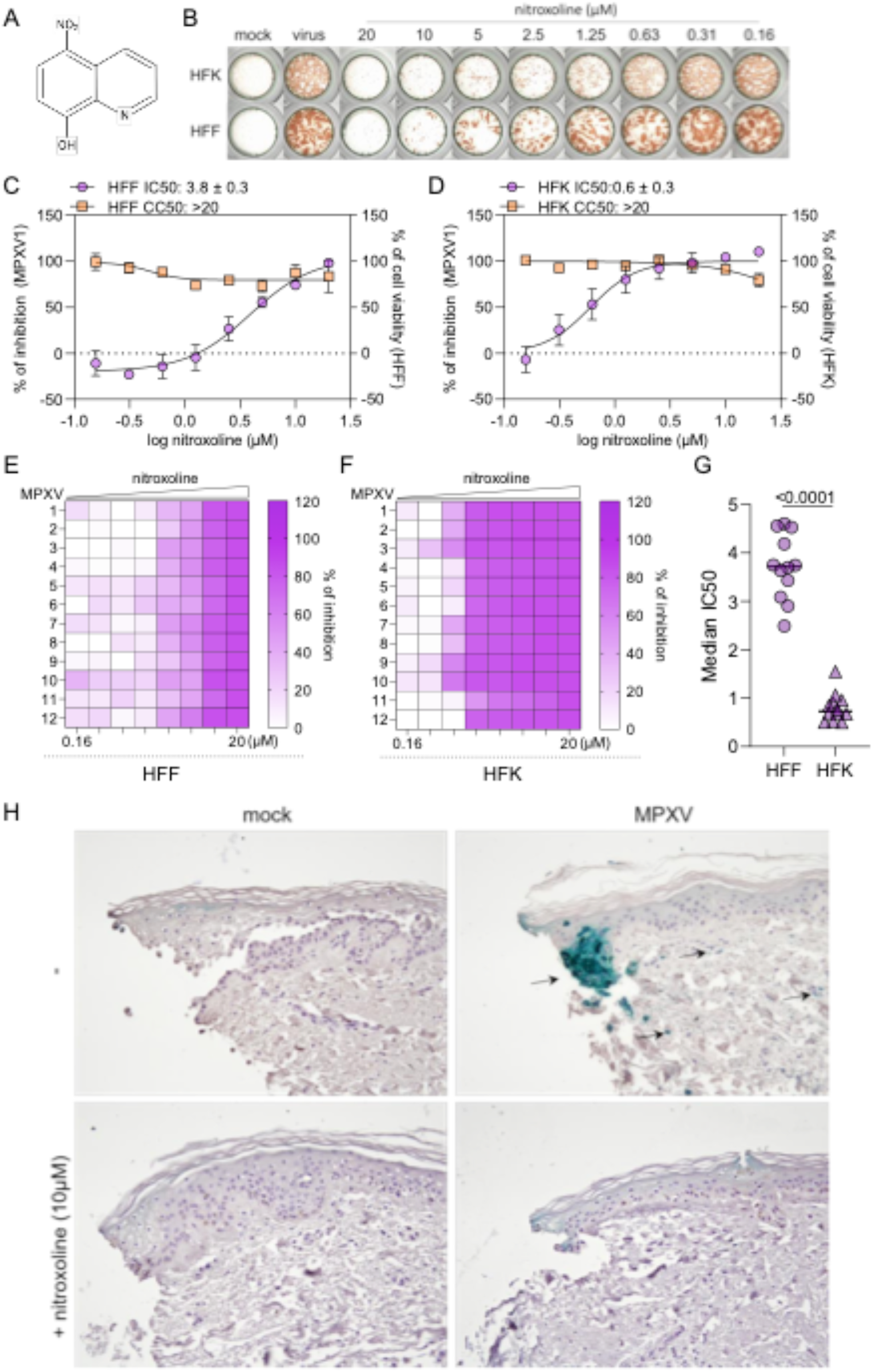
Effects of nitroxoline on mpox virus (MPXV) replication in primary human fibroblasts (HFF), keratinocytes (HFK), and a skin explant model. A) Chemical structure of nitroxoline. B-D) Concentration-dependent effects of nitroxoline on mpox virus isolate 1 (MPXV1, MOI 0.01) infection in HFF and HFK, as indicated by immunostaining. IC50 = concentration that inhibits mpox virus infection by 50% as indicated by immunostaining; CC50 = concentration that reduces cell viability by 50% as indicated by MTT assay. E,F) Concentration-dependent effects of nitroxoline on HFF and HFK infection with 12 mpox virus isolates, as indicated by immunostaining. G) Nitroxoline IC50s in HFF and HFK. H) Effects of nitroxoline on MPXV1 infection in a skin explant model. Primary human skin tissue was infected with 10^6^ TCID50/ml of MPXV1 per well in 500 μL with or without nitroxoline treatment at 10 μM for 48h. Then, the skin tissue was embedded into paraffin and sectioned. Virus infection was detected by immunohistochemical staining.

When added to the culture medium together with the virus, nitroxoline inhibited mpox virus infection in HFF and HFK in a dose-dependent manner (Figure 1B-D) as indicated by immunostaining. The nitroxoline concentrations that reduced virus immunostaining by 50% (IC50) ranged from 2.4 to 4.6 μM in HFF and from 0.5 to 1.5 μM in HFK (Figure 1E-G, Suppl. Table 1). Nitroxoline did not affect cell viability in the tested concentration range of up to 20μM (Figure 1C,D). Notably, nitroxoline may interfere with different orthopoxviruses, as it also inhibited vaccinia virus infection at a similar IC50 (5.2μM) as mpox virus infection (Suppl. Figure 1).

Time-of-addition experiments (Suppl. Figure 2A) showed that nitroxoline interferes with the mpox virus replication cycle post viral entry (Suppl. Figure 2B,C). Nitroxoline inhibited mpox virus infection in a similar way when it was added two hours post infection (Suppl. Figure 2B,C) as when it was added simultaneously with the virus (Figure 1C,D). However, nitroxoline addition together with virus followed by a washing step after a two-hour entry period was not effective (Suppl. Figure 2B,C). Moreover, nitroxoline only reduced virus titres (as determined by PCR for genomic mpox virus DNA), when added after the two-hour virus absorption period, but not when it was present only during the entry period (Suppl. Figure 2D).

To investigate the antiviral effects of nitroxoline in the context of the skin architecture, we used primary human split-thickness skin grafts that preserve the histology and complexity of the skin [Hendriks et al., 2021]. Skin grafts were infected with 10^6^ TCID50/mL of mpox virus isolate 1 (MPXV1), and the infection was visualised by immunohistochemical staining for virus antigen after 48h. As depicted in Figure 1H, pronounced infection was detected in the epidermis. Moreover, clusters of infected cells or single infected cells were located in the dermis (Figure 1H). These findings are in line with the known patterns of mpox infection in human skin [Stagles et al., 1985; Reed et al., 2004]. Nitroxoline (10μM) treatment strongly reduced the number of mpox-infected cells.

### Effects of nitroxoline analogues on mpox virus infection

Next, we investigated a set of nine nitroxoline analogues for anti-mpox virus activity in HFF (Figure 2A). Only compounds 1 (IC50: 1.8 ± 0.3μM), 7 (IC50: 3.6 ± 1.5μM), and 9 (IC50: 2.1 ± 0.1μM) displayed a similar antiviral activity as nitroxoline (IC50: 2.1 ± 0.7μM) (Figure 2B). The active nitroxoline analogues all harboured halogen ions at positions 5 and 8 and a hydroxy group at position 9. Notably, compound 9 is clioquinol, another antibiotic that is clinically being used for the treatment of different skin infections [Wykowski et al., 2022] (Figure 2). Further research will have to show whether it may be possible to identify nitroxoline analogues with a higher anti-mpox virus activity than nitroxoline.

**Figure 2.**
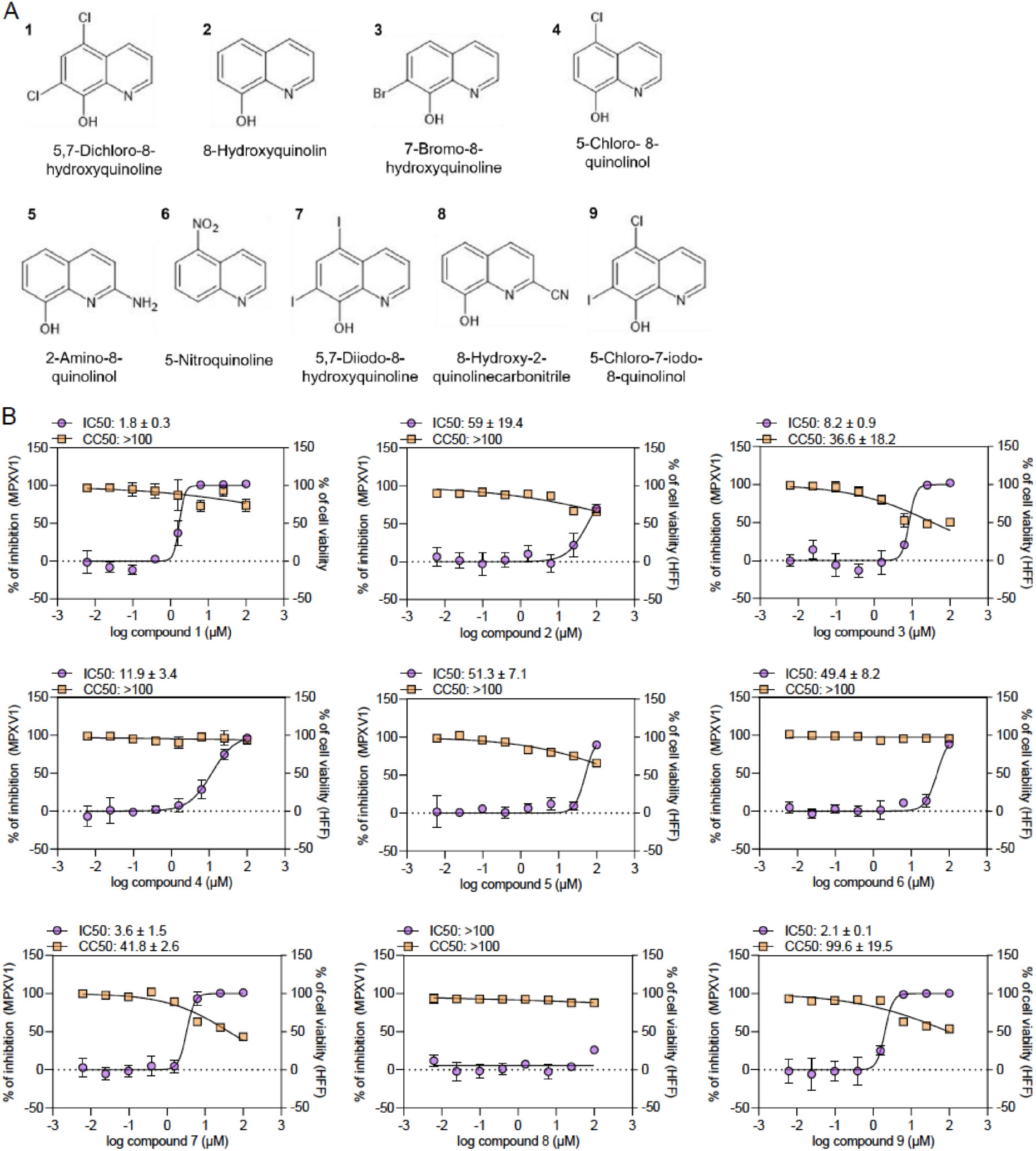
Effects of nitroxoline analogues on mpox virus infection. A) Chemical structures of the investigated nitroxoline analogues. B) Dose-response curves indicating compound effects on mpox virus (MPXV1) infection as indicated by immunostaining of MOI 0.01-infected primary human foreskin fibroblasts (HFF) and MTT assay in mock-infected HFF.

### Nitroxoline interferes with mpox virus-induced cellular signalling pathways

Nitroxoline inhibits bacterial growth by chelating cations that are required by bacterial metalloenzymes, and the addition of cations such as Mg^2+^ and Mn^2+^ abrogates its antibacterial activity [Repac Antić et al., 2022]. In contrast, the addition of Mg^2+^, Mn^2+^, or other divalent cations did not affect the antiviral activity of nitroxoline (Figure 3A,B) indicating a different mode of antiviral action.

**Figure 3.**
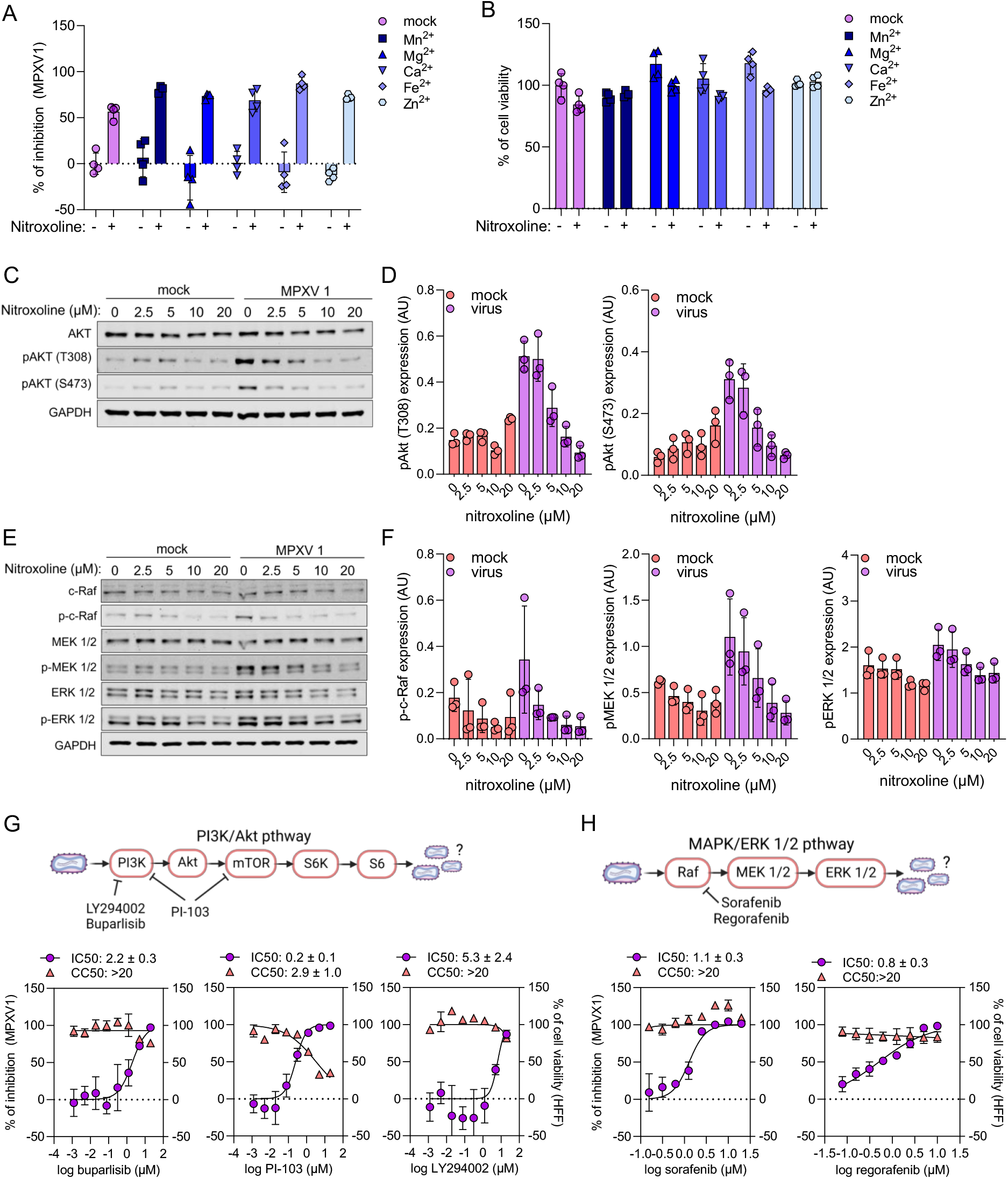
Nitroxoline interferes with mpox virus-induced cellular signalling pathways. A,B) Cations known to inhibit antibacterial effects of the chelator nitroxoline did not inhibit nitroxoline’s antiviral activity as indicated by immunostaining in mpox virus isolate 1 (MPVX1) MOI 0.01-infected primary human foreskin fibroblasts (HFF, A) and did not affect cell viability in the presence of nitroxoline as indicated by MTT assay in mock-infected HFF (B). C,D) Nitroxoline reduces AKT phosphorylation in a dose-dependent manner as indicated by Western blot. E,F) Nitroxoline reduces Raf, MEK, and ERK phosphorylation in a dose-dependent manner as indicated by Western blot. G,H) PI3K, PI3K/mTOR, and Raf inhibitors suppress mpox virus infection in a dose-dependent manner, as determined in MPVX1 MOI 0.01-infected HFF. Compound effects on cell viability were detected by MTT assay in mock-infected HFF.

However, nitroxoline inhibited virus-induced PI3K/AKT signalling (as indicated by AKT phosphorylation, Figure 3C,D) and MAPK signalling (as indicated by RAF, MEK, and ERK phosphorylation, Figure 3E,F) in a dose-dependent manner. Moreover, inhibitors of PI3K/AKT/mTOR (buparlisib, LY294002, PI-103) and MAPK (sorafenib, regorafenib) signalling inhibited mpox virus infection (Figure 3G,H). These data agree with previous findings showing that nitroxoline inhibits PI3K/AKT/mTOR and Raf/MEK/ERK signalling [Chang et al., 2015; Xu et al., 2019; Palicelli et al., 2021] and that orthopoxvirus replication critically depends on PI3K/AKT/mTOR and Raf/MEK/ERK signalling [Kindrachuk et al., 2012; Beerli et al., 2019; Peng et al., 2020]. Taken together, these data suggest that nitroxoline inhibits mpox virus infection at least in part by interference with these two host cell signalling pathways.

### Nitroxoline inhibits a tecovirimat-resistant mpox virus strain

Based on experience with other antiviral drugs, there is concern that tecovirimat-resistant viruses may emerge [DeLaurentis et al., 2022; Gessain et al., 2022]. Hence, we established a tecovirimat-resistant mpox virus strain (Figure 4A). ARPE cells were infected with mpox virus isolate 1 (MPXV1) at a multiplicity of infection (MOI) of 0.01 in the presence of tecovirimat 4μM. After seven days, medium was removed and replaced by fresh tecovirimat 4μM-containing medium. After a total incubation time of 14 days, cytopathogenic effects were detected, and the tecovirimat-resistant substrain was expanded (Figure 4A).

**Figure 4.**
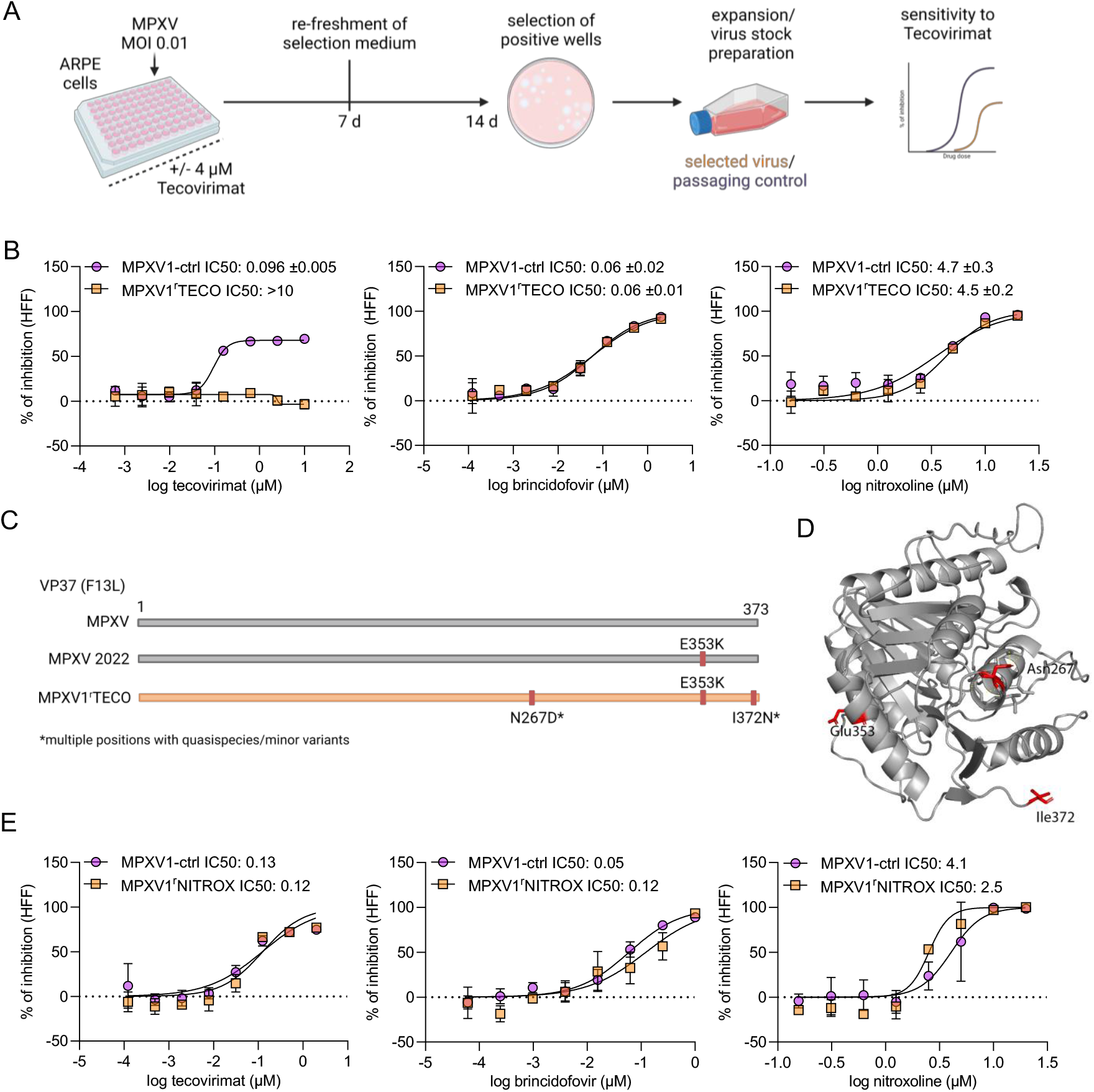
Effects of nitroxoline and brincidofovir on a tecovirimat-adapted mpox virus strain. A) Scheme of the one-round adaptation approach for the generation of a tecovirimat-resistant sub-strain (MPXV1^r^TECO)) by exposure of the mpox virus isolate 1 (MPXV1) to tecovirimat 4μM. B) Dose-dependent effects and IC50 values of tecovirimat, brincidofovir, and nitroxoline in primary human foreskin fibroblasts infected with MPXV1 or MPXV1^r^TECO at an MOI 0.01 as detected by immunostaining. C) Amino acid sequence changes in F13L (the target of tecovirimat) from mpox viruses from the current global outbreak (MPXV 2022) including MPXV1 and MPXV1^r^TECO relative to pre-outbreak sequences. N267D and I372N were previously shown to mediate tecovirimat resistance [Duraffour et al., 2015; FDA, 2022]. D) Location of E (Glu, glutamate) 353, N (Asn, asparagine) 267 and I (Ile, isoleucine) in the F13L structure. The change from N (Asn, asparagine) to D (Asp, aspartate) in position 267 results in the loss forms hydrogen bonds with E263. E) Sensitivity of a MPXV1 substrain that was cultivated for three passages in the presence of nitroxoline (5μM) to tecovirimat, brincidofovir, and nitroxoline as indicated by immunostaining 48h post infection with MOI 0.01.

The resulting tecovirimat-selected MPXV1 sub-strain (MPXV1^r^TECO) displayed a pronounced tecovirimat resistance as indicated by an IC50 of >10μM compared to an IC50 of 0.096μM of a passaging control (Figure 4B). Whole genome virus sequencing indicated three amino acid sequence changes (E353K, N267D, I372N) in F13L (TP37, gp45), the target of tecovirimat (Figure 4C, Figure 4D, Suppl. Table 2). E353K is shared between isolates from the current global outbreak and was shown not to affect tecovirimat efficacy [Bojkova et al., 2022]. In contrast, N267D and I372N were previously shown to provide resistance to tecovirimat and are, hence, likely responsible for the observed tecovirimat resistance [Duraffour et al., 2015; FDA, 2022]. Notably, MPXV1^r^TECO remained sensitive to both brincidofovir and nitroxoline (Figure 4B).

In contrast to MPXV1 cultivation in the presence of tecovirimat, MPXV1 cultivation in the presence of nitroxoline (5μM) did not result in reduced virus sensitivity to nitroxoline, tecovirimat, or brincidofovir, although the incubation time was increased to three passages (Figure 4E).

### Effects of nitroxoline on *E. coli, N. gonorrhoeae*, and herpes viruses

Next, we evaluated the activity of nitroxoline against bacterial (*Escherichia coli, Neisseria gonorrhoeae*) and viral (varicella zoster virus, herpes simplex virus type 1) pathogens that are commonly co-transmitted with mpox viruses [Hughes et al., 2020; Girometti et al., 2022; Patel et al., 2022].

14 *E. coli* patient isolates displayed nitroxoline sensitivity as indicated by disk diffusion (inhibition zones: 17-24mm), agar dilution (maximum inhibitory concentrations (MICs): 4-8μg/ml corresponding to 21-42 μM), and applying clinical breakpoints set by EUCAST (Suppl. Figure 3A, Suppl. Table 3). Susceptibility testing of *N. gonorrhoeae* revealed similar results (inhibition zones: 22-25mm and MICs of 4-8 μg/ml), indicating a susceptible phenotype (Suppl. Figure 3B, Suppl. Table 4).

In contrast to mpox virus infection, nitroxoline inhibited varicella zoster virus and herpes simplex virus type 1 infection only at a concentration of 20μM (Suppl. Figure 3C, Suppl. Figure 3D).

### Combination of nitroxoline with antiviral drugs

Antiviral combination therapies can result in increased efficacy and reduced resistance formation [White et al., 2021]. In agreement, brincidofovir and tecovirimat displayed increased antiviral activity when used in combination against different orthopoxviruses in preclinical model systems [Quenelle et al., 2007; Chen et al., 2011]. In this context, nitroxoline displayed additive activity in combination with tecovirimat and synergistic activity in combination with brincidofovir against mpox virus infection, as determined by the method of Chou & Talalay [Chou, 2006] (Figure 5).

**Figure 5.**
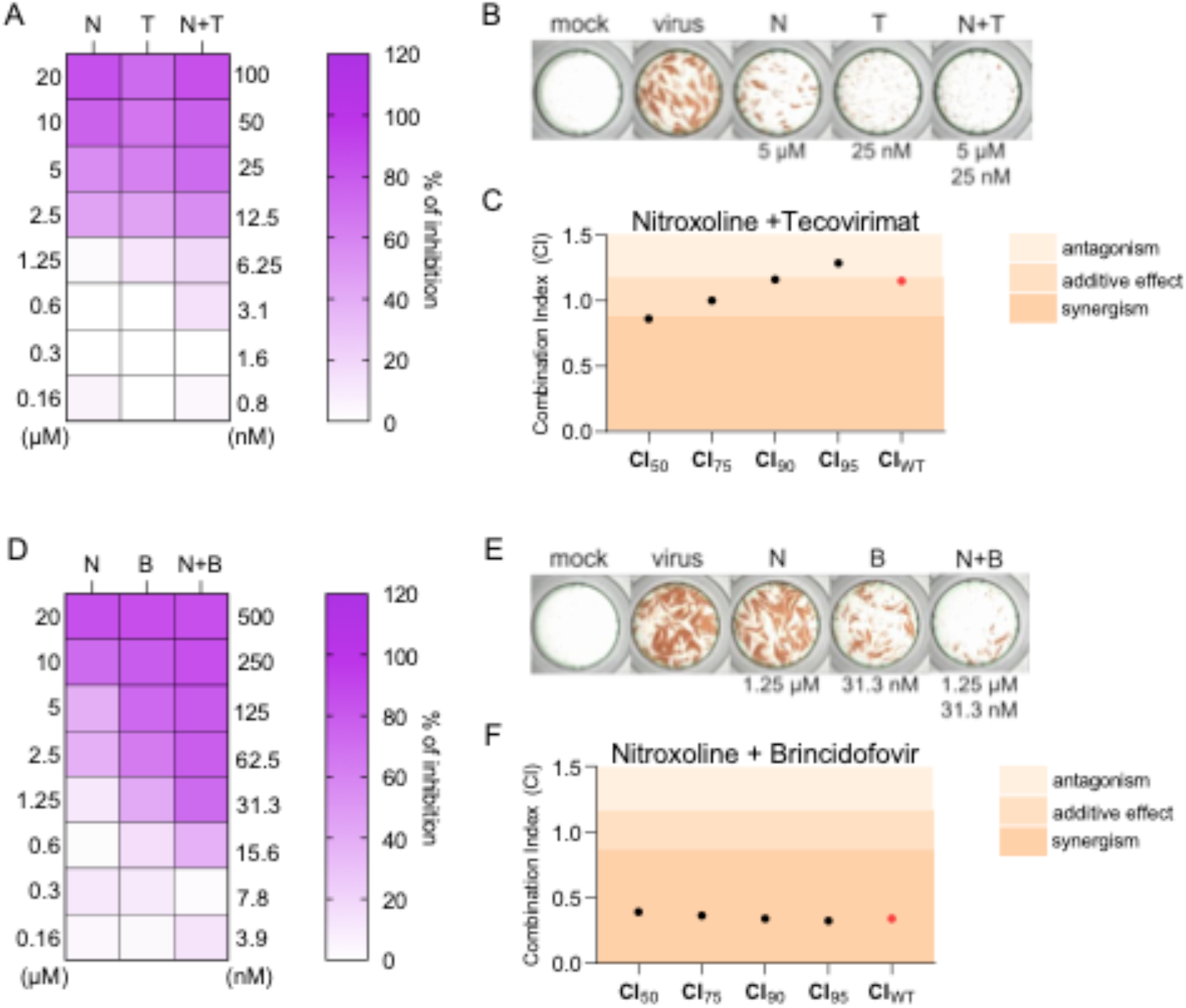
Antiviral activity of nitroxoline in combination with tecovirimat and brincidofovir. A) Dose-dependent effects of nitroxoline (N, 0.16-20μM), tecovirimat (T, 0.8-100μM), and their combination in primary human foreskin fibroblasts (HFF) infected with mpox virus isolate 1 (MOI 0.01) as indicated by immunostaining. B) Representative immunostaining images illustrating the combined effects of nitroxoline (N) and tecovirimat (T). C) Determination of the combination index (CI) of nitroxoline (N) and tecovirimat (T) following the method of Chou and Talalay [Chou, 2006]. D) Dose-dependent effects of nitroxoline (N, 0.16-20μM), brincidofovir (B, 3.9-500μM), and their combination in HFF infected with mpox virus isolate 1 (MOI 0.01) as indicated by immunostaining. E) Representative immunostaining images illustrating the combined effects of nitroxoline (N) and brincidofovir (B). F) Determination of the CI of nitroxoline (N) and brincidofovir (B) following the method of Chou and Talalay [Chou, 2006].

## Discussion

Nitroxoline is an FDA-approved antibiotic that has been used for more than 50 years for the treatment of acute and recurrent urinary tract infections. It is currently used as a first-line therapy for uncomplicated urinary tract infections in Germany due to its excellent activity towards both Gram-negative bacteria and fungi as well as its favourable safety profile [Naber et al., 2014; Wijma et al., 2018]. In this study, nitroxoline effectively inhibited the replication of 12 mpox virus isolates from the current outbreak. The nitroxoline IC50s (0.5 - 4.6μM) were within the range of therapeutic plasma levels that have been reported to reach between 30 and 50μM [Wijma et al., 2018]. Moreover, nitroxoline also suppressed mpox virus replication in a skin explant model. The investigation of nine nitroxoline analogues did not identify a compound with superior activity against mpox virus relative to nitroxoline.

Tecovirimat (F13L inhibitor) and brincidofovir (DNA polymerase inhibitor) are the antiviral drugs that are currently mainly considered for mpox treatment [DeLaurentis et al., 2022; Gessain et al., 2022; Huang et al., 2022; Bojkova et al., 2022]. There are concerns about the potential emergence of tecovirimat-resistant mpox virus strains [DeLaurentis et al., 2022; Gessain et al., 2022], and the formation of a tecovirimat-resistant vaccinia virus was described in an immunocompromised acute myeloid leukaemia patient after inoculation with the vaccinia virus-based ACAM2000 smallpox vaccine [Lederman et al., 2012].

We established a tecovirimat-resistant mpox virus strain (MPXV1^r^TECO), which harboured the known tecovirimat resistance mutations N267D and I372N, by adapting mpox virus isolate 1 (MPXV1) to tecovirimat in a one round selection step using a high tecovirimat concentration (4μM). This approach is similar to that previously described for the generation of a tecovirimat-resistant cowpox virus [Yang et al., 2005]. In contrast, another study reported the establishment of tecovirimat-resistant poxviruses by exposure to step-wise increasing drug concentrations to be a lengthy process (6-18 months) that is not always successful [Duraffour et al., 2015]. The reasons underlying these discrepancies remain unclear. It may be possible that the currently circulating mpox viruses harbour small tecovirimat-resistant subpopulations that become readily selected and enriched in response to tecovirimat treatment.

Notably, MPXV1^r^TECO remained sensitive to nitroxoline (and brincidofovir). In contrast to tecovirimat, nitroxoline treatment of mpox virus using the same approach did not result in the formation of a nitroxoline-resistant strain. This agrees with evidence suggesting that the targeting of host cell factors by antiviral drugs is associated with reduced resistance formation compared to agents that directly target virus proteins [De Clercq, 2002; Zheng et al., 2022].

Moreover, nitroxoline exerted additive antiviral effects in combination with tecovirimat and synergistic effects in combination with brincidofovir. Hence, its clinical anti-mpox virus activity in humans can be tested in combination with these antivirals without depriving study participants of these more established options. Additionally, nitroxoline combination therapies with increased activity may delay resistance formation by monkey pox virus [White et al., 2022].

Nitroxoline was previously reported to inhibit a genetically modified Japanese encephalitis virus strain in the hepatoma cell line Huh7 [Zhang et al., 2020], but information on its antiviral mechanisms of action is lacking. Nitroxoline exerts its antibacterial effects by chelating metal ions including Fe2^+^, Mn2^+^, and Mg2^+^ [Pelletier et al., 1995]. Although poxviruses depend on the availability of bivalent cations for effective replication [Li et al., 2016; Xu J et al., 2019], the antiviral activity of nitroxoline was not affected by the addition of metal ions. This shows that nitroxoline’s antiviral and antibacterial mechanisms of action differ substantially.

Our further research demonstrated that nitroxoline inhibits mpox virus replication at least in part by interfering with the PI3K/AKT/mTOR and Raf/MEK/ERK host cell signalling pathways that are critical for orthopoxvirus replication [Kindrachuk et al., 2012; Beerli et al., 2019; Peng et al., 2020]. Notably, the clinically approved Raf inhibitors sorafenib and regorafenib also suppressed mpox virus infection at nontoxic concentrations.

In agreement with previous findings [Pelletier et al., 1995; Naber et al., 2014; Fuchs et al., 2019], nitroxoline was also effective against *N. gonorrhoeae* and *E.coli*, two sexually transmitted bacteria that are commonly co-transmitted with mpox virus in the current outbreak [Girometti et al., 2022; Patel et al., 2022]. Moreover, nitroxoline inhibited infection caused by herpes simplex virus type 1 and varicella zoster virus, two herpes viruses that are often detected together with mpox virus [Hughes et al., 2020; Girometti et al., 2022; Patel et al., 2022], albeit at higher concentrations (>10μM) than those blocking mpox virus infection. These effects may also be caused by inhibition of PI3K/AKT/mTOR and Raf/MEK/ERK signalling, as interference with these signalling pathways has also been described to affect herpes virus replication [Rahaus et al., 2007; Tiwari & Shukla, 2010; Seo et al., 2015; Lesch et al., 2019; Madavaraju et al., 2021].

In conclusion, nitroxoline inhibited mpox viruses from the current global outbreak, including a tecovirimat-adapted strain, at therapeutically achievable concentrations. Moreover, it increased the activity of and can be used in combination with the two approved anti-poxvirus drugs tecovirimat and brincidofovir. Nitroxoline is potentially also a readily available alternative to these antivirals, as the use of brincidofovir is associated with significant adverse effects and tecovirimat stocks are insufficient to cover the current outbreak [Adler et al., 2022; Gessain et al., 2022; Johri et al., 2022; Pfäfflin et al., 2022]. Finally, nitroxoline is also effective against pathogens that are co-transmitted with mpox virus in the current outbreak, such as sexually transmitted bacterial and viral illnesses [Girometti et al., 2022; Patel et al., 2022]. Thus, nitroxoline is a repurposing candidate for the treatment of mpox virus that may also have potential for the treatment of neglected mpox disease in endemic areas in Africa and for the control and ideally prevention of future global outbreaks [Alakunle & Okeke, 2022].

## Methods

### Cell culture

Human foreskin fibroblasts (HFF) and human foreskin keratinocytes (HFK) were isolated as previously described [Zöller et al., 2014; Wilhelm et al., 2021] according to the Declaration of Helsinki principles and in agreement with the institutional review board (112/06; 386/14). HFF were cultured in Dulbecco’s Modified Eagle Medium (DMEM) with 4.5g/ml glucose supplemented with 5% foetal bovine serum (FBS) and 100IU/ml penicillin. HKF were cultured in DermaLife K (CellSystems) supplemented with 100IU/ml penicillin. The cell lines ARPE (ATCC) and HaCaT (CLS Cell Lines Service) were cultured in minimal essential medium (MEM) supplemented with 10% FBS, 100 IU/mL penicillin, and 100 μg/mL streptomycin. All cell lines were regularly authenticated by short tandem repeat (STR) analysis and tested for mycoplasma contamination.

### Mpox virus isolation and production

Mpox virus clinical isolates were obtained by culturing swabs from the patient’s lesions on Vero cells. After appearance of cytopathogenic effect (CPE) both cells and supernatant were frozen at −80°C. For virus stock preparation, the human keratinocyte cell line HaCaT was utilised. Briefly, cells were incubated with 50μL of infectious inoculum for 72h and subsequently frozen at −80°C until further processing. After thawing, supernatants were centrifuged at 150g for 10min and virus stocks stored at −80°C. Virus titres were determined as TCID50/mL using confluent HFF in 96-well microtiter plates.

### Antiviral assay

Confluent cells in 96-well plates were infected with mpox virus isolates at MOI 0.01 and incubated at 37°C for 48h. Drug inhibitory effects were determined by immunocytochemistry staining of mpox virus. Briefly, cells were fixed with acetone:methanol (40:60) solution and immunostaining was performed using an anti-Vaccinia Virus antibody (1:4000 dilution, #ab35219 Abcam, Berlin, Germany), which was detected with a peroxidase-conjugated anti-rabbit secondary antibody (1:1,000, Dianova), followed by addition of AEC substrate. The mpox virus positive area was scanned and quantified by the Bioreader^®^ 7000-F-Z-I microplate reader (Biosys). The results are expressed as percentage of inhibition relative to virus control which received no drug.

### Cell viability assay

Cell viability was measured by the 3-(4,5-dimethylthiazol-2-yl)-2,5-diphenyltetrazolium bromide (MTT) dye reduction assay 96-well plates. 25 μL of MTT solution (2 mg/mL in PBS) were added per well, and the plates were incubated at 37 °C for 4 h. After this, the cells were lysed using 100 μL of a buffer containing 20% sodium dodecylsulfate and 50% *N*,*N*-dimethylformamide with the pH adjusted to 4.7 at 37 °C for 4 h. Absorbance was determined at 560 nm (reference wavelength 620 nm) using a Tecan infinite M200 microplate reader (TECAN).

### Mpox virus isolate assignment to clades

Total DNA from viral stocks was isolated using the QIAamp DNA Blood Kit (Qiagen, Hilden, Germany) according to the manufacturer’s instructions. DNA was subjected to qRT-PCR analysis using the Luna Universal qPCR Master Mix Protocol (New England Biolabs, Frankfurt am Main, Germany) and a CFX96 Real-Time System, C1000 Touch Thermal Cycler (Bio-Rad, Feldkirchen, Germany). Primers detecting mpox virus were adapted from Liu et al. 2010 [Li et al., 2010].

### Split-thickness skin model

Skin samples derived from surplus split skin not used for wound cover were placed in PBS and perforated by microneedle pre-treatment (Segminismart^®^, Nicosia, Cyprus) to facilitate virus infection as described [Tajpara et al., 2019]. Then, 3×3mm skin pieces were infected with 10^6^ TCID50/mL of mpox virus isolate 1 (MPXV1) per well in 500 μL with or without nitroxoline (10 μM). 48 h post infection, tissue samples were formalin-fixed, paraffin-embedded (FFPE), and cut into 4μm sections. After deparaffinisation and heat-induced epitope retrieval (Target Retrieval Solution pH9, Agilent-Dako, S2367, Santa Clara, U.S.A.), sections were incubated with a primary anti-vaccinia virus antibody (1:10.000, Abcam, ab35219, Berlin, Germany), followed by incubation with secondary anti-rabbit IgG-horseradish peroxidase conjugates (ZytoChem HRP Kit, HRP-125, Zytomed Systems, Berlin, Germany), and visualisation using HistoGreen (Histo Green Kit, Linaris, LIN-E109, Frankfurt Germany) as peroxidase substrate. All experiments were performed according to the Declaration of Helsinki principles and in agreement with the institutional review board (112/06; 386/14)

### Immunoblot analysis

Whole-cell lysates were prepared using Triton-X sample buffer containing protease inhibitor cocktail (Roche). The protein concentration was assessed by using DC Protein assay reagent (Bio-Rad Laboratories). Equal protein loads were separated by sodium dodecyl sulfate-polyacrylamide gel electrophoresis and proteins were transferred to nitrocellulose membranes (Thermo Scientific). For protein detection the following primary antibodies were used at the indicated dilutions: AKT (Cell Signaling, #9272, 1:1000), phospho-AKT T308 (Cell Signaling, #2965, 1:1000), phospho-AKT S473 (Cell Signaling, #4060, 1:1000), c-Raf (Cell Signaling, #9422, 1:1000), phospho-c-Raf S338 (Cell Signaling, #9327, 1:1000), ERK1/2 (Acris, #AP00033P4-N, 1:1000), phospho-ERK1/2 T202/Y204 (Cell Signaling, #9106, 1:1000), GAPDH (Cell Signaling, #2118, 1:4000), MEK1/2 (Cell Signaling 1:1000, #9122, 1:1000), phospho-MEK1/2 S217/221 (Cell Signaling, #9121, 1:1000). Protein bands were visualized using IRDye-labeled secondary antibodies at dilution 1:40000 (LI-COR Biotechnology, IRDye^®^800CW Goat anti-Rabbit, #926-32211 and IRDye^®^800CW Goat anti-Mouse IgG, #926-32210) and Odyssey Infrared Imaging System (LI-COR Biosciences).

### Drug combination assay

To evaluate antiviral activity of nitroxoline in a combination with tecovirimat and brincidofovir, the compounds were applied alone or in fixed combinations at 1:2 dilutions using HFF monolayers. Subsequently the cells were infected with MPVX 1 at MOI 0.01 for 48 h. The calculation of IC_50_, IC_75_, IC_90_ and IC_95_ for single drugs and their combinations as well as combination indexes (CIs) was performed using the software CalcuSyn (Biosoft) based on the method of Chou and Talalay [Chou, 2006]. The weighted average CI value (CI_wt_) was calculated according to the formula: CI_wt_ [CI_50_ + 2CI_75_ + 3CI_90_ + 4CI_95_]/10. CI_wt_ values were calculated for mutually exclusive interactions where CI_wt_ <0.8 indicates synergism, CI_wt_ between 0.8-1.2 indicates additive effects, and CI_wt_ >1.2 suggest antagonism.

### Selection of Tecovorimat-resistant variant

ARPE cells were seeded in 96-well plate 48 h prior infection and treatment. The cells were treated with 4 μM of Tecovirimat and subsequently infected with MPXV1 at MOI 0.01. Untreated cells were used as passaging control. After 7 days the selection medium containing 4 μM of Tecovirimat was refreshed and the cells were incubated for additional 7 days. The positive wells displaying plaques were harvested and expanded to viral stocks. The resistance development was validated in antiviral assay.

### Complete virus genome sequencing

Up to 5ng extracted DNA were used for library preparation using the KAPA Hyper Prep Kit (Roche) according to manufacturer’s instructions. Resulting libraries were quantified on a TapeStation System (Agilent), equimolar pooled, and paired-end sequenced on an Illumina MiniSeq sequencer (Illumina, 300 cycles). Reads were mapped against ON563414.2 using Geneious Prime v2022.0.1 and manually curated.

### Effect of nitroxoline on HSV-1 and VZV

Antiviral efficacy of nitroxoline against two sexually transmitted herpesviruses, HSV-1 and VZV, was evaluated in HFF and ARPE cells, respectively. Briefly, confluent layers of HFF or ARPE cells were treated with nitroxoline and infected with HSV-1 McIntyre strain (ATCC) at MOI 0.01 for 24 h or with VZV clinical isolate [Schmidt-Chanasit et al., 2008] at MOI 0.1 for 48 h. Subsequently, the cell were fixed with acetone:methanol (40:60) solution and immunostained with antibody directed against HSV-1 (#ab9533, Abcam, Berlin, Germany) or against VZV (IE62-specific mAb, Chemicon, Billerica, MA, USA), which was detected with a peroxidase-conjugated anti-rabbit or anti-mouse secondary antibody (1:1,000, Dianova), respectively, followed by addition of AEC substrate. The virus positive area was quantified by the Bioreader^®^ 7000-F-Z-I microplate reader (Biosys). The results are expressed as percentage of inhibition relative to non-treated virus control.

### Bacterial isolates and antibiotic susceptibility testing

All bacterial isolates were recovered from patients hospitalized at the Goethe University Hospital in Frankfurt. Reference strains *Escherichia coli* ATCC 25922 and *Neisseria gonorrhoeae* ATCC 49226 were obtained from DSMZ (German Collection of Microorganisms and Cell Culture, Braunschweig, Germany).

Antimicrobial susceptibility was determined by disc diffusion (Liofilchem^®^, Roseto degli Abruzzi, Italy) using Mueller Hinton agar (Oxoid™, Thermo Fisher, Darmstadt, Germany) for *E. coli* and Chocolate agar with Vitox (Oxoid™, Thermo Fisher, Darmstadt, Germany) for *N. gonorrhoeae*. Agar dilution was performed with Mueller Hinton agar for *Escherichia coli* and GC agar supplemented with hemoglobin solution and BBL™ IsoVitaleX™ (Becton, Dickinson and Company, Le Pont de Claix, France) for *Neisseria gonorrhoeae* with increasing concentrations of nitroxoline. Additionally, broth microdilution was performed with cation-adjusted Mueller-Hinton broth for *E. coli*.

Inhibition zones and minimum inhibitory concentrations (MICs) were evaluated and interpreted according to EUCAST guidelines for *E. coli* due to the undefined criteria for *N. gonorrhoeae* [https://www.eucast.org/clinical_breakpoints/].

### Structural Modelling

The mpox F13L protein structure was modelling using Phyre2 [Kelley et al., 2015] (with default settings). Phyre2 generated a high confidence model for 93% of the protein sequence.

### Statistics

The results are expressed as the mean ± standard deviation (SD) of the number of biological replicates indicated in figure legends. The statistical significance is depicted directly in graphs and the statistical test used for calculation of p values is indicated in figure legends. GraphPad Prism 9 was used to determine IC50 values.

## Acknowledgement

We thank Kerstin Euler, Sebastian Grothe, Daniel Janisch and Lena Stegmann for their technical support.

