## Supplementary material for "Repurposing of the antibiotic nitroxoline for the treatment of mpox": Suppl. Figures and Tables

Suppl Fig 1

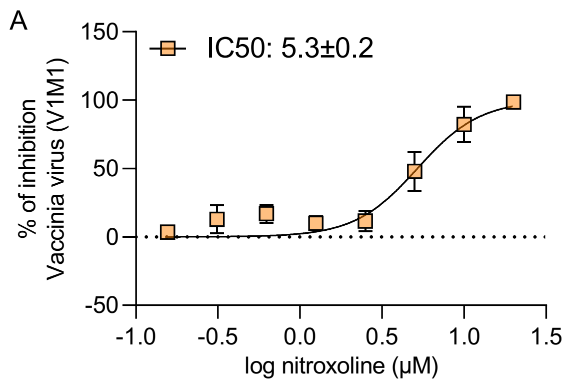


**Suppl. Figure 1. Effects of nitroxoline on vaccinia virus (V1M1) replication in primary human fibroblasts (HFF).** Concentration-dependent effects of nitroxoline on V1M1 virus infection in HFF, as indicated by immunostaining. IC50 = concentration that inhibits V1M1 virus infection by 50% as indicated by immunostaining.

Suppl Fig 2

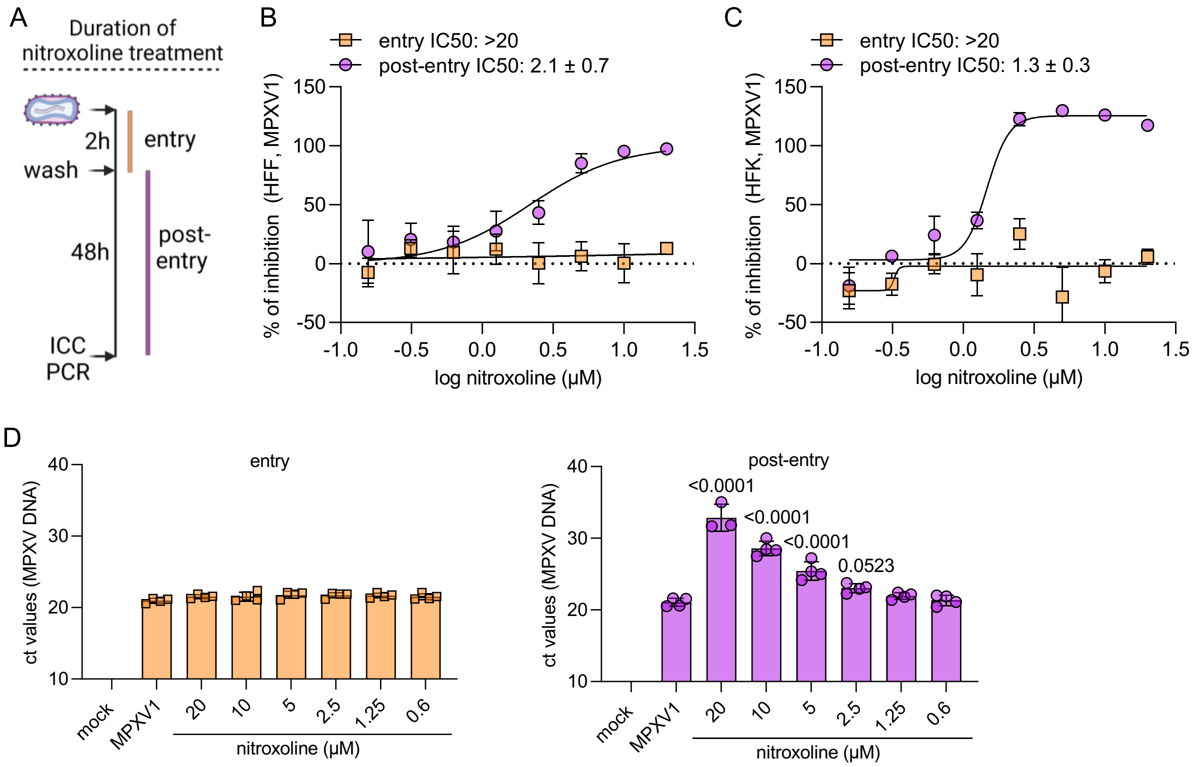


**Suppl. Figure 2. Effects of nitroxoline on mpox virus (MPVX) replication in primary human fibroblasts (HFF) and keratinocytes (HFK).** A) Design of time-of-addition experiments. B-D) Effects of nitroxoline on mpox virus (MPXV1) infection and replication in HFFs when exclusively administered during (entry) or after (post-entry) the 2h virus adsorption period as indicated by immunostaining (B,C) or qPCR for genomic mpox virus DNA (D).

Suppl Figure 3

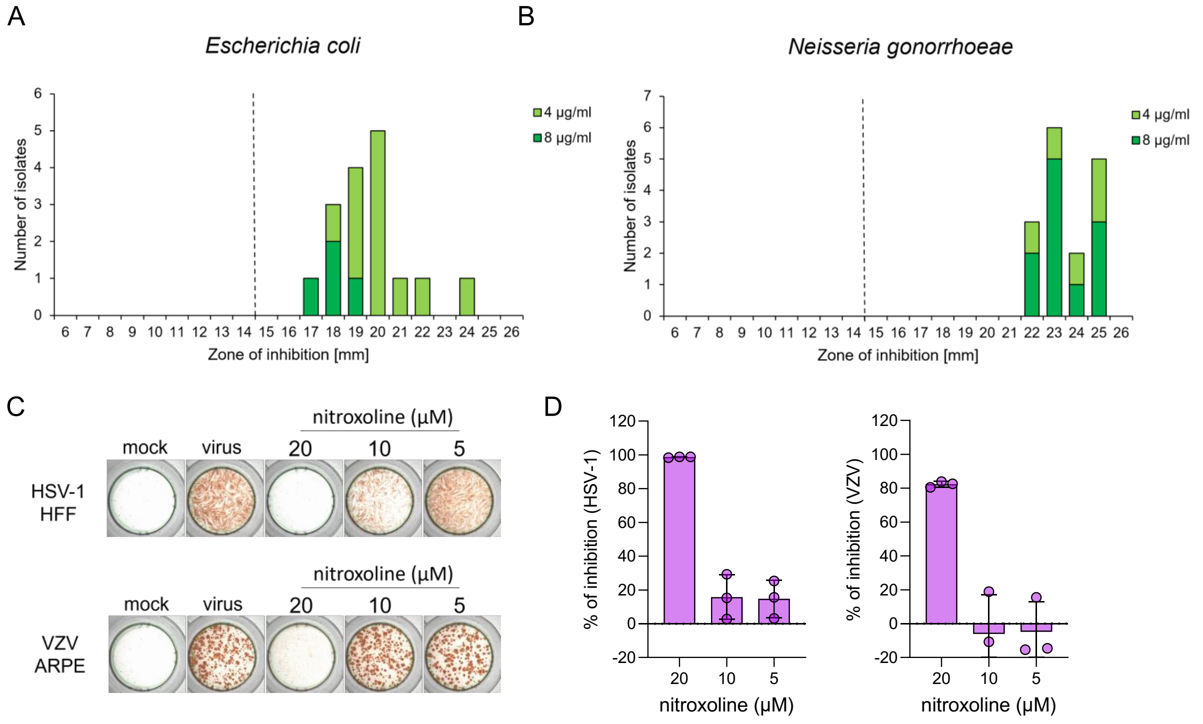


**Suppl. Figure 3. Effects of nitroxoline on pathogens that are commonly transmitted together with mpox virus.** Antimicrobial susceptibility to nitroxoline was determined by disc diffusion for *Escherichia coli (A)* and *Neisseria gonorrhoeae* (B). C,D) Effects of nitroxoline against herpes simplex virus type 1 (HSV-1) and Varicella zoster virus (VZV). Effects on HSV-1 were determined in McIntyre strain (MOI 0.01)-infected human foreskin fibroblasts by immunostaining 24h post infection. Effects on VZV were detected in ARPE cells infected with a clinical isolate [Schmidt-Chanasit et al., 2008] at MOI 0.1 by immunostaining 48h post infection.

**Suppl. Table 1. Nitroxoline concentrations that inhibit infection of primary human foreskin fibroblasts (HFF) and keratinocytes (HFK) with clinical mpox virus (MPXV) isolates (MOI 0.01) by 50% (IC50) as indicated by immunostaining 48h post infection.**

| Mpox virus isolate | HFF[1] | HFK[1] |
|---|---|---|
| MPXV1 | 4.19 ± 1.24 | 0.68 ± 0.12 |
| MPXV2 | 4.60 ± 0.12 | 0.69 ± 0.09 |
| MPXV3 | 3.75 ± 0.51 | 0.50 ± 0.25 |
| MPXV4 | 3.74 ± 0.71 | 0.91 ± 0.06 |
| MPXV5 | 3.70 ± 0.05 | 0.85 ± 0.03 |
| MPXV6 | 3.43 ± 0.74 | 0.96 ± 0.09 |
| MPXV7 | 3.09 ± 0.81 | 0.69 ± 0.05 |
| MPXV8 | 3.64 ± 0.39 | 0.76 ± 0.04 |
| MPXV9 | 4.56 ± 1.28 | 0.52 ± 0.03 |
| MPXV10 | 2.49 ± 0.67 | 0.50 ± 0.08 |
| MPXV11 | 4.53 ± 0.59 | 1.55 ± 0.01 |
| MPXV12 | 2.90 ± 0.61 | 1.07 ± 0.13 |

[1] mean ± S.D.

**Suppl. Table 2. Amino acid substitutions in the tecovirimat-adapted MPXV1 substrain MPXV1$^{r}$TECO as determined by complete virus genome sequencing relative to a reference genome (ON563414.2).** Sequence changes in MPXV1$^{r}$TECO relative to MPXV1 are highlighted in bold.

| ID Frankfurt | Total reads | Reads mapped against MPX [a] | Mean genome coverage | Coding sequence coverage with >10 reads | Amino acid substitutions (compared to outbreak reference sequence [a]) |
|---|---|---|---|---|---|
| MPXV1 | 10,096,514 | 3,247,443 | 2470.9 | 100% [b] | gp10: 913 nt deletion (affecting AA221-525, new stop codon);<br>gp157: D4N |
| MPXV1$^{r}$TECO | 16,421,220 | 5,874,779 | 4462.0 | 100% [b] | gp10: 913 nt deletion (affecting AA221-525, new stop codon); gp44: **D549N**; gp45: **N267D**, **I372N**; gp157: D4N; **gp161:S306L** |

[a] ON563414.2 - Monkeypox virus isolate MPXV_USA_2022_MA001, 2022

[b] 913 nt deletion in gp010

**Suppl. Table 3.** Effects of nitroxoline on *Escherichia coli* determined using clinical isolates and the reference strain ATCC 25922 by disc diffusion and agar dilution assay.

| Method | | Disc diffusion | Agar dilution | Broth microdilution |
|---|---|---|---|---|
| | | Inhibition zone (mm) | MIC (µM) | MIC (µM) |
| E.coli (isolates) | 2652 | 19 | 21 | 21 |
| | 2966 | 22 | 21 | 21 |
| | 3513 | 20 | 21 | 21 |
| | 3719 | 20 | 21 | 21 |
| | 3726 | 17 | 42 | 42 |
| | 4041 | 20 | 21 | 21 |
| | 4559 | 20 | 21 | 21 |
| | 4954 | 18 | 42 | 21 |
| | 5173 | 17 | 42 | 42 |
| | 5301 | 19 | 21 | 21 |
| | 5752 | 19 | 21 | 21 |
| | 5762 | 18 | 21 | 21 |
| | 5763 | 16 | 42 | 42 |
| | 7975 | 21 | 21 | 21 |
| | 8667 | 20 | 21 | 21 |
| | ATCC 25922 | 24 | 21 | 10.5 |

**Suppl. Table 4.** Effects of nitroxoline on *Neisseria gonorrhoeae* determined using clinical isolates and the reference strain ATCC ATCC49226 by disc diffusion and agar dilution assay.

| Method | | Disc diffusion | Agar dilution |
|---|---|---|---|
| | | Inhibition zone (mm) | MIC [µM] |
| N. gonorrhoeae (isolate) | 428 | 25 | 42 |
| | 429 | 24 | 42 |
| | 432 | 22 | 42 |
| | 436 | 25 | 42 |
| | 439 | 23 | 42 |
| | 440 | 25 | 42 |
| | 441 | 23 | 42 |
| | 442 | 23 | 42 |
| | 444 | 23 | 42 |
| | 520 | 25 | 21 |
| | 550 | 23 | 42 |
| | 570 | 23 | 21 |
| | 600 | 25 | 21 |
| | 690 | 24 | 21 |
| | 740 | 22 | 21 |
| | ATCC49226 | 22 | 42 |
